# First-described recently discovered non-toxic vegetal-derived furocoumarin preclinical efficacy against SARS-CoV-2: a promising antiviral herbal drug

**DOI:** 10.1101/2020.12.04.410340

**Authors:** Iván José Galindo-Cardiel, Adriana Toledo Núñez, María Celaya Fernández, Ariel Ramírez Labrada, Iratxe Uranga-Murillo, Maykel Arias Cabrero, Julian Pardo, Ezio Panzeri

**Affiliations:** WorldPathol Global United S.A. (WGUSA), Zaragoza, Spain; ICE-P Life S.L., Barcelona, Spain; Unidad de Nanotoxicología e Inmunotoxicología (UNATI), Instituto de Investigación Sanitaria de Aragón (IISA), Zaragoza, Spain; Immunotherapy, Inflammation and Cancer, Instituto de Investigación Sanitaria Aragón (IISA), Zaragoza, Spain; Instituto de Carboquímica, CSIC, Zaragoza, Spain; Fundación ARAID, Zaragoza, Spain; Dpto. Microbiologia, Medicina Preventiva y Salud Pública, Fac. Medicina, Universidad de Zaragoza, Spain

**Keywords:** *Angelica archangelica*, antiviral, COVID19, herbal drug, SARS-CoV-2, therapeutic

## Abstract

Severe acute respiratory syndrome coronavirus 2 (SARS-CoV-2) is the aetiology of coronavirus disease 2019 (COVID19) pandemic. ICEP4 purified compound (ICEP4) is a recently discovered furocoumarin-related purified compound derived from the roots and seeds of *Angelica archangelica* (herbal drug). ICEP4-related herbal preparations have been extensively used as active herbal ingredients in traditional medicine treatments in several European countries. Extraction method of patent pending ICEP4 (patent application no. GB2017123.7) has previously shown strong manufacturing robustness, long-lasting stability, and repeated chemical consistency. Here we show that ICEP4 presents a significant *in vitro* cytoprotective effect in highly virulent-SARS-CoV-2 challenged Vero E6 cellular cultures, using doses of 34.5 and 69 μM. No dose-related ICEP4 toxicity was observed in Vero E6 cells, M0 macrophages, B, CD4+ T and CD8+ T lymphocytes, Natural Killer (NK) or Natural Killer T (NKT) cells. No dose-related ICEP4 inflammatory response was observed in M0 macrophages quantified by IL6 and TNFα release in cell supernatant. No decrease in survival rate was observed after either 24 hr acute or 21-day chronic exposure in *in vivo* toxicity studies performed in *C. elegans*. Therefore, ICEP4 toxicological profile has demonstrated marked differences compared to others vegetal furocoumarins. Successful ICEP4 doses against SARS-CoV-2-challenged cells are within the maximum threshold of toxicity concern (TTC) of furocoumarins as herbal preparation, stated by European Medicines Agency (EMA). The characteristic chemical compounding of ICEP4, along with its safe TTC, allow us to assume that the first-observation of a natural antiviral compound has occurred. The potential druggability of a new synthetic ICEP4-related compound remains to be elucidated. However, well-established historical use of ICEP4-related compounds as herbal preparations may point towards an already-safe, widely extended remedy, which may be ready-to-go for large-scale clinical trials under the EMA emergency regulatory pathway. To the best of the authors’ knowledge, ICEP4-related herbal drug can be postulated as a promising therapeutic treatment for COVID19.

## 1. Introduction

Severe acute respiratory syndrome coronavirus 2 (SARS-CoV-2) is an enveloped non-segmented positive-sense single-stranded RNA virus (genus Betacoronavirus, subfamily *Orthocoronavirinae*). SARS-CoV-2 is the aetiology of the coronavirus disease 2019 (COVID19) pandemic (1). Social distancing, case identification, contact tracing, quarantine and isolation are postulated as the main strategies to reduce viral spreading. Despite worldwide research efforts and some very promising advances, no effective antiviral drugs, or mitigant sanitary products against SARS-CoV-2 infection currently exist, therefore, current pharmacological therapy is mostly restricted towards mitigating the associated symptoms (2, 3).

There is continuous interest in searching for alternative antiviral drugs among phytochemical extracts, medicinal plants, and aromatic herbs. Discovery and production of novel antiviral drugs frequently occurs from spices, herbal medicines, essential oils (EOs), and distilled natural products (4). Coumarins comprise a large class of compounds found within medicine herbal preparations (5–7). Coumarins are found at high levels in some EOs, particularly cinnamon bark oil, cassia leaf oil, and lavender oil. Coumarin is also found in fruits (e.g. bilberry, cloudberry), green tea, and other foods, such as chicory (8). Most coumarins occur in higher plants, with the richest sources being the *Rutaceae* and *Umbelliferae*. Although distributed throughout all parts of the plant, the coumarins occur at the highest levels in the fruits, followed by the roots, stems, and then leaves. Environmental conditions and seasonal changes may influence the occurrence of coumarins in various parts of the plant (9). Psoralens are natural products that are linear furanocoumarins (most furanocoumarins can be regarded as derivatives of psoralen or angelicin), which are extremely toxic to a wide variety of prokaryotic and eukaryotic organisms. Some important psoralen derivatives are xanthotoxin, imperatorin, bergapten and nodekenetin (8, 9). The demonstrated activities of coumarins include anticoagulant, anticancer, antioxidant, antiviral, anti-diabetic, anti-inflammatory, antibacterial, antifungal, and anti-neurodegerative properties as drugs, as well as the ability to act as fluorescent sensors for biological systems (10).

The genus *Angelica litoralis* is comprised of over 90 species spread throughout most areas of the globe (11). More than half of these species are used in traditional therapies, while some of them are included in several national and European pharmacopoeias (12–15). Bioactive constituents in different *Angelica* species include coumarins, EOs, polysaccharides, organic acids and acetylenic compounds (16). *In vitro* testing confirmed cytotoxic (17, 18), anti-inflammatory (19), antibacterial (20), antifungal (21), neuroprotective (22) and serotonergic (23) activities for extracts obtained from a range of *Angelica* species.

Reducing viral replication at the beginning of SARS-CoV-2 infection and, subsequently, the associated degree of immunopathological damage, is a critical step to mitigate and cure COVID19 (2). ICEP4 (patent pending, application n° GB2017123.7) is an *Angelica archangelica*-based purified compound with previous evidence of antiviral and oncolytic *in vitro* effects (ICE-P Life, data not shown). ICEP4-related herbal preparations have been extensively used as active herbal ingredients in traditional medicine treatments in several countries, including the EU and US (12–15). The main objective of this work was to evaluate the possible cytoprotective effects of the ICEP4 extract against SARS-CoV-2 challenge by means of Crystal Violet staining, a technique used as an indirect quantification method for cell death. In parallel, the potential cytotoxic effects of ICEP4 were assessed, both *in vitro* and *in vivo*, using standard EMA-accepted methods. Both objectives should support ICEP4 use as a new, safe antiviral herbal drug for COVID19 treatment that acts via stopping viral spreading at targeted-SARS-CoV-2 epithelium, without compromising the host immune response.

## 2. Material and Methods

### 2.1. Study design

Efficacy assays were performed in biosafety level 3 (BSL3) facilities at Zaragoza (Spain) (WGUSA, laboratory reference 747735/2014). Cytotoxic studies and replication of *in vitro* efficacy studies were independently repeated in biosafety level 2 (BSL2) facilities at UNATI (IISA, Zaragoza, Spain) and BSL3 facilities at the University of Zaragoza in order to demonstrate experimental repeatability and inter-laboratory consistency of obtained results.

ICEP4 efficacy against SARS-CoV-2 challenge was evaluated *in vitro* using the Crystal Violet staining technique. Cellular viability testing was performed by measuring the percentage of stained cells per well after SARS-CoV-2 challenge was carried out in triplicate, in three different tests. Three doses of ICEP4 were tested, along with the proper negative/positive controls, in three different tests (T1, T2, and T3).

The *in vitro* toxicity of ICEP4 at various concentrations, was analysed in immune cells. C57BL/6 (*Mus musculus*) (B6)-mouse-derived bone marrow monocytes (BMC) were differentiated to M0 macrophages and incubated for 24 hrs with the various doses of ICEP4. Proper ICEP4-negative and highest concentration-used diluent (SHAM) controls were included for comparison. After 24 hrs, cell viability was determined using the PrestoBlue^TM^ assay. Additionally, the macrophage inflammatory response was determined by measuring IL6 and TNFα expression in cell supernatants. On the other hand, B6-mouse-derived splenocytes were incubated for 24 hrs with ICEP4, along with appropiate controls as indicated for macrophages. Subsequently, cells were labelled with CD3-FITC, CD8-APC, CD4-VioBlue and NK1.1-APCVio770 or CD19-PE, CD3-FITC together with Annexin-V-PE or -APC and dead cells within the T, NK and NKT or B cells populations were evaluated via flow cytometry (GALLIOS, Beckman Coulter).

*In vivo* acute and chronic toxicity was evaluated using the survival rate of ICEP4-challenged synchronised cultured glp-4 mutant *Caenorhabditis elegans*. Both assays were carried out at 25°C in three separate experiments; the duration of the assays were 24 hrs and 21-days for acute and chronic toxicity, respectively.

### 2.2. Raw Material

#### 2.2.1. ICEP4 plant-derived extract dosing

Five mg of original 9-years-old certified-batch ICEP4 was submitted by Mr. Ezio Panzeri for examination. ICEP4 extract comes from seeds and roots (herbal drug) of *A. archangelica*. Briefly, 700 g of coarsely comminated plant-derived material was successively extracted for 36 hrs with 15 L of methanol in a Soxhlet extractor (Quickfit^TM^ large-scale extractor IIEX). The extracts were concentrated under reduced pressure (rotary evaporator 9200/1). ICEP4 purification was performed with high-performance liquid chromatography to reach 98% purity (UPLC High-performance Liquid Chromatographer XEVO TQD, Waters, US), showing a characteristic consistent chemical profile after a second ICEP4 manufactured batch (WGUSA, data not shown) (Acquity UPLC PDA QDA <ESI-MS>, Waters, US) (Figure 1). The standard dose for toxicology and efficacy studies of this first described natural-derived compound were determined considering the toxicologic data from previously described *Angelica*-related furocoumarin (CAS number 66-97-7) (24). The most appropriate solvent was taken from previous chemical characterization studies (WGUSA, data not shown).

**Figure 1.**
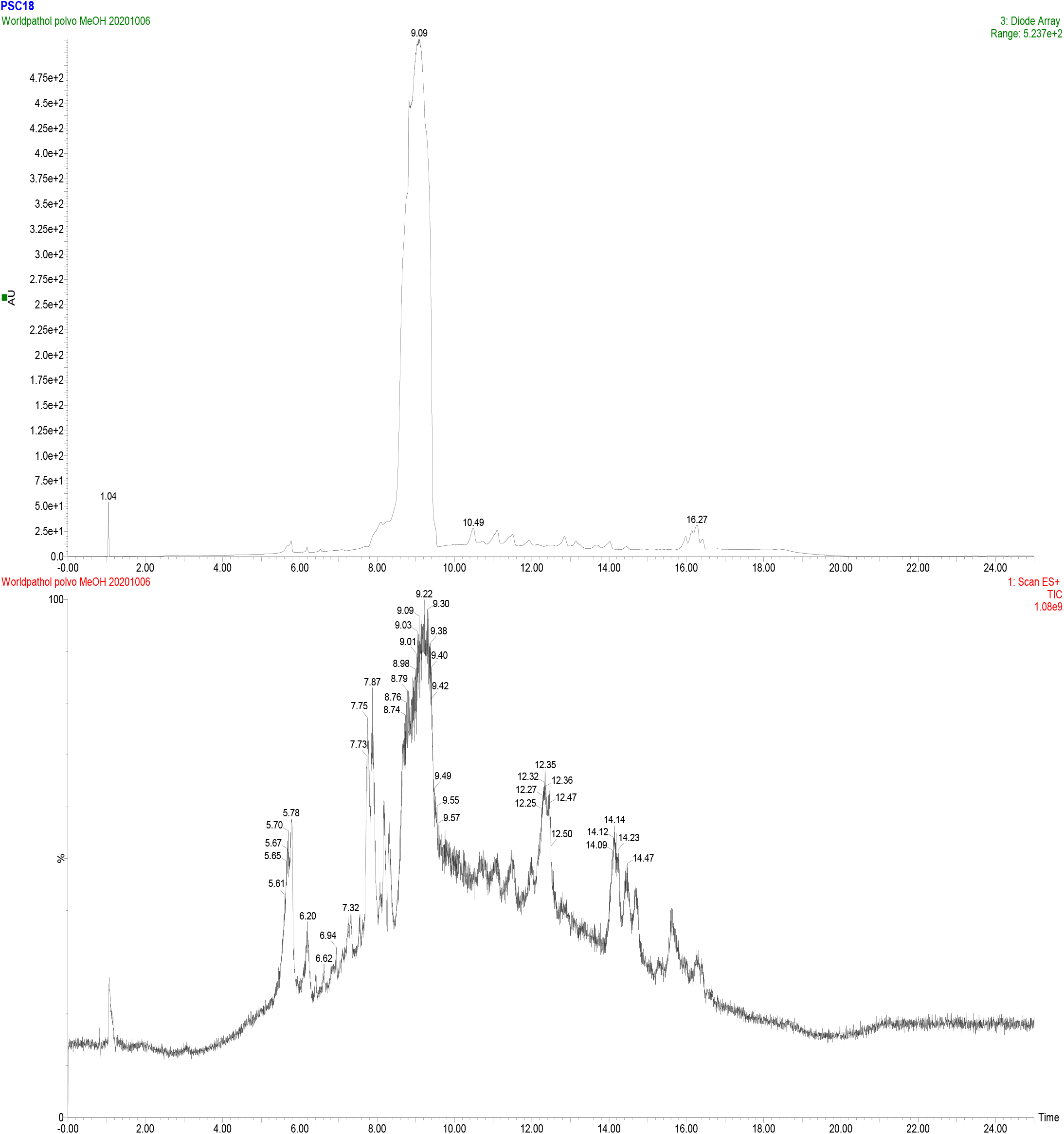
Units of absorbance of the original 9-years-old ICEP4 batch. Methanol was used as solvent. Note that there is saturation in expected wavelength. 320 nm at a retention time of 9 minutes (Acquity UPLC PDA QDA <ESI-MS>, Waters, US).

The starting point was a stock dilution of original ICEP4 in ethanol with a concentration of 1 mg/mL (4.6 mM). Raw material was diluted at concentrations between 1μM and 1mM to assess preliminary ICEP4 toxicity in macrophages and immune cells. Higher concentration-used solvent was used as SHAM control of cellular viability.

Cellular viability was assessed as an indicator of ICEP4 efficacy following SARS-CoV-2 challenge. Three doses of ICEP4 were selected based on previous immunotoxicology data. The threshold of toxicological concern (TTC) for furocoumarins is 1.5 μg/Kg of daily exposure, therefore, *in vitro* experiments were carried out taking these values into account to calculate the maximum concentration (24). Cytotoxicity assays were performed in 96-well plates with 6.9 μM, 34.5 μM, and 69 μM concentrations of the ICEP4 maximum dose. ICEP4 doses were vehiculated, among others, with ethanol 0.15%, 0.75% and 1.5%. The highest concentration-used solvent was included as a SHAM control for cellular viability.

For *in vivo* studies, a 1 mM dose of ICEP4 was added to a 25° C-cultivated infertile strain of *C. elegans* for acute and chronic studies (10.000 times higher than reference dose).

#### 2.2.2. SARS-CoV-2 strain

A high-pathogenic strain of SARS-CoV-2 was isolated and cultured from a 72-year old patient at University Clinical Hospital Lozano Blesa (Zaragoza, Spain). Second-passage vials with the SARS-CoV-2 strain were provided by Dr. Julian Pardos (IISA, UNATI, Zaragoza, Spain). Virus was maintained and cultured following UNATI protocols in BSL3 facilities at Zaragoza (WGUSA, Spain). The tissue culture infectious dose 50% (TCID50) was determined to be 1.47 x 10^6^/mL. The same strain and TCID50 was used at UNATI facilities for repeatability and inter-laboratory consistency studies.

### 2.3. Methods

#### 2.3.1. SARS-CoV-2-challenged ICEP4-treated efficacy assay

Vero E6 cells were provided by Eugenia Puentes (Biofabri, Porriño, Spain) and cultured following provider’s descriptions. Cellular cultures were maintained at a density of 10^5^ cell/mL in Vero E6 10% FBS (Sigma F7524) Dulbecco's Modified Eagle Medium (Lonza, Ref BE12-614F) throughout the study, at 37 °C with 5% CO_2_ and 90% humidity. The efficacy assays were performed in 96-well plates (Nunclon Delta Surface 167008 Thermo) with a density of 10^4^ cells/well. Vero E6 cells were seeded a day prior to viral infection. The experiment was carried out following the design described in Table 1. Briefly, the plate contained internal growing controls of Vero E6 cells, the described concentrations of ethanol (vehicle-related toxicity control), and ICEP4 doses (both compound toxicity control and cytoprotective effect). SARS-CoV-2 was added after 1 hr of incubation (37 °C), and the plates were then incubated for 72 hrs at 37 °C and 5% CO_2_ following viral challenge.

**Table 1.** Descriptive statistics of *in vitro* challenge test. ICEP4 doses are 6.9 (WP1), 34.5 (WP2) and 69 (WP3) μM per well. Different ethanol concentrations (0.15%, 0.50%, 0.75%) were added as SHAM control. TCID50 is considered positive control and the main infective group to be compared with treatment. TCID50-SARS-CoV-2 10-fold is considered maximum infective positive internal control (cell death). Treatment-negative Control (CT) group was composed by non-treated Vero E6 cells in 2% FBS medium (maximum cell viability).

|  | *CT* | *Et_0,15* | *Et_0,5* | *Et_0,75* | *WP1* | *WP2* | *WP2* | *TCID50* | *SARS+*<br>*WP1* | *SARS+*<br>*WP2* | *SARS+*<br>*WP3* | *SARS*<br>*10-f* |
| --- | --- | --- | --- | --- | --- | --- | --- | --- | --- | --- | --- | --- |
| Mean | 8 | 8 | 7.33 | 7.67 | 7.33 | 7.33 | 7.67 | 5 | 7.00 | 7.67 | 7.67 | 0 |
| Standard error | 0 | 0 | 0.33 | 0.33 | 0.33 | 0.33 | 0.33 | 1 | 0.58 | 0.33 | 0.33 | 0 |
| Median | 8 | 8 | 7 | 8 | 7 | 7 | 8 | 4 | 7 | 8 | 8 | 0 |
| Mode | 8 | 8 | 7 | 8 | 7 | 7 | 8 | 4 | 7 | 8 | 8 | 0 |
| Standard deviation | 0 | 0 | 0.58 | 0.58 | 0.58 | 0.58 | 0.58 | 1.73 | 1 | 0.58 | 0.58 | 0 |
| Variance | 0 | 0 | 0.33 | 0.33 | 0.33 | 0.33 | 0.33 | 3 | 1 | 0.33 | 0.33 | 0 |

Cellular viability was observed by Crystal Violet staining. Briefly, 72 hrs after SARS-CoV-2 challenge, cells were fixed with 4% paraformaldehyde (Panreac 252931.1212) for 1 hr at room temperature. Cells were then stained with Crystal Violet solution (0.5% crystal violet and 20% methanol) (Sigma, C0775) (Panreac 131091.1212). Cellular viability was directly observed by inverted microscope (DM IL LED Leica). Strong positive cellular staining (blue) was considered as an indicator of viable cells (more than 75% of the well stained). Intermediate or weak cellular staining was considered to represent as unviable cells (less than 75% of the well stained). Counting was performed by two different technologists per experiment.

#### 2.3.2. *In vitro* cytotoxicity assays

##### 2.3.2.1. M0 macrophage differentiation from mouse bone marrow-derived cells

Bone marrow derived cells (BMDCs) were obtained from a minimal-disease certified mouse (C57BL/6 -*M. musculus*- -B6-, Charles River, US). Femurs and tibias were dissected from the euthanised mouse and, under sterile conditions, the bone marrow was eluted by injecting DMEM or RPMI medium through the bone cavity. Erythrocytes were lysed and final BMDC suspension was adjusted to 10^6^ cells/mL. BMDMs were differentiated into M0 macrophages after an incubation period of 6 days with BMDM medium, and, finally, seeded at a concentration of 5·10^4^ cells/well in 96-well plates. After 24 hrs, ICEP4 was dosed by making 10-fold serial dilutions and the cells were incubated for an additional 24 hrs.

##### 2.3.2.2. Isolation of mouse splenocytes

A minimal-disease certified mouse (B6, Charles River, US) was killed by cervical dislocation. The spleen was then carefully extracted and mashed through a cell strainer. Splenocytes were washed with RPMI and centrifuged at 1200 rpm for 5 min. Splenocytes were counted and adjusted to 10^6^ cells/mL.

##### 2.3.2.3. Macrophage cellular viability by PrestoBlue^TM^ assay and inflammatory response

Cell viability was analysed by PrestoBlue^TM^ HS (high sensitivity) assay following the manufacturer’s instructions. PrestoBlue^TM^ HS contain resazurin and a propriety buffering system (#P50200, ThermoFisher, US). Absorbance was measured using an iMark™ Microplate Absorbance Reader (BioRad, Germany). Activation of the inflammatory response in macrophages was analysed quantifying the cytokines IL6 and TNFα in cell supernatants by ELISA (Ready-Set-Go kit, eBiosciences) following the manufacturer’s instructions.

##### 2.3.2.4. Lymphocytes analysis by flow cytometry

Splenocytes were incubated with the same concentrations of the ICEP4 molecule used for macrophage studies. After 24 hrs, the cells were collected and washed, and cell viability was analysed by annexin V staining in T, B, NK, and NKT cells, as indicated in 2.1.

#### 2.3.3. *In vivo* toxicity: *C. elegans* assay

##### 2.3.3.1. *C. elegans* strain

The *C. elegans* strain used was the glp-4 mutant. *Caenorhabditis elegans* gene glp-4 was identified by the temperature-sensitive allele bn2 where mutants raised at the restrictive temperature (25 °C) produce adults that are essentially germ cell deficient *C. elegans*.

##### 2.3.3.2. Culturing and synchronization

*C. elegans* worms were propagated on nematode growth media (NGM) agar plates with kanamycin 50 μg/mL and streptomycin 100 μg/mL at 20 °C (NGM Lite, US Biological Life Sciences, Swampscott in Massachusetts, US) using *E. coli OP50* as a source of food. Due to the presence of worms at different developmental stages in cultures, a synchronization process consisting of killing larvae and adult worms using a combination of NaOH and NaClO was followed, as described elsewhere (25). Eggs obtained from synchronization were resuspended and plated on a NGM agar plate without *E. coli OP50* (ISSA, Zaragoza, Spain), in order to allow the eggs to hatch and reduce developmental differences in new larvae due to differences in egg ages. *E. coli OP50* was added to the NGM agar plate 24 hrs later.

##### 2.3.3.3. Acute survival assay

L1 larvae obtained from synchronization were cultured at 25 °C until worms developed to L4 stage. L4 worms were harvested from plates and washed 3 times with M9. Approximately 15 worms per well were placed in a 96-well flat bottom microtiter plate and treated with different ICEP4 doses at 25 °C. A total of 45 worms (3 wells) were assessed for each dose of ICEP4. Worms without treatment served as negative controls. Three different independent survival assays were carried out.

##### 2.3.3.4. Chronic survival assay

Worms were cultured as previously described and treated with different ICEP4 doses. Worms without treatment served as negative controls. Survival assays were carried out for 21 days at 25 °C. Every 7 days, ICEP4 and *E. coli* OP50 were added to the *C. elegans*. Chronic toxicity worms were seeded and counted twice a week calculating the percentage of worms that survived with respect to the number of worms at time zero. Three independent experiments were performed.

### 2.4. Statistics of efficacy studies

Efficacy data were analysed by Microsoft^®^ Excel^®^ STATS (Microsoft 365 MSO - 16.0.13231.20110-32 bits, ID 00265-80196-36405-AA936). Results were presented as Mean ± SD (Standard Deviation). One-way ANOVA was used to confirm statistical differences among multiple groups between treated and non-treated groups. ICEP4 – TCID50 groups were analysed by two-sample t-test, assuming equal variances, to confirm significant differences. Significant differences are indicated by: *P < 0.05; **P < 0.01; and ***P < 0.001. P < 0.05 was considered as significant. Results from UNATI were analysed together with WGUSA obtained data, in order to check robustness and repeatability.

## 3. Results

### 3.1. SARS-CoV-2-challenged ICEP4-treated efficacy assay

Descriptive statistics including UNATI results are shown in Table 1. Maximum SD was observed in the TCID50 group, mostly due to outlier results in the first replication of the experiment (Figure 2). After this first trial, more coherent and consistent TCID50 results (5±1,73) were obtained across the rest of the replications.

**Figure 2.**
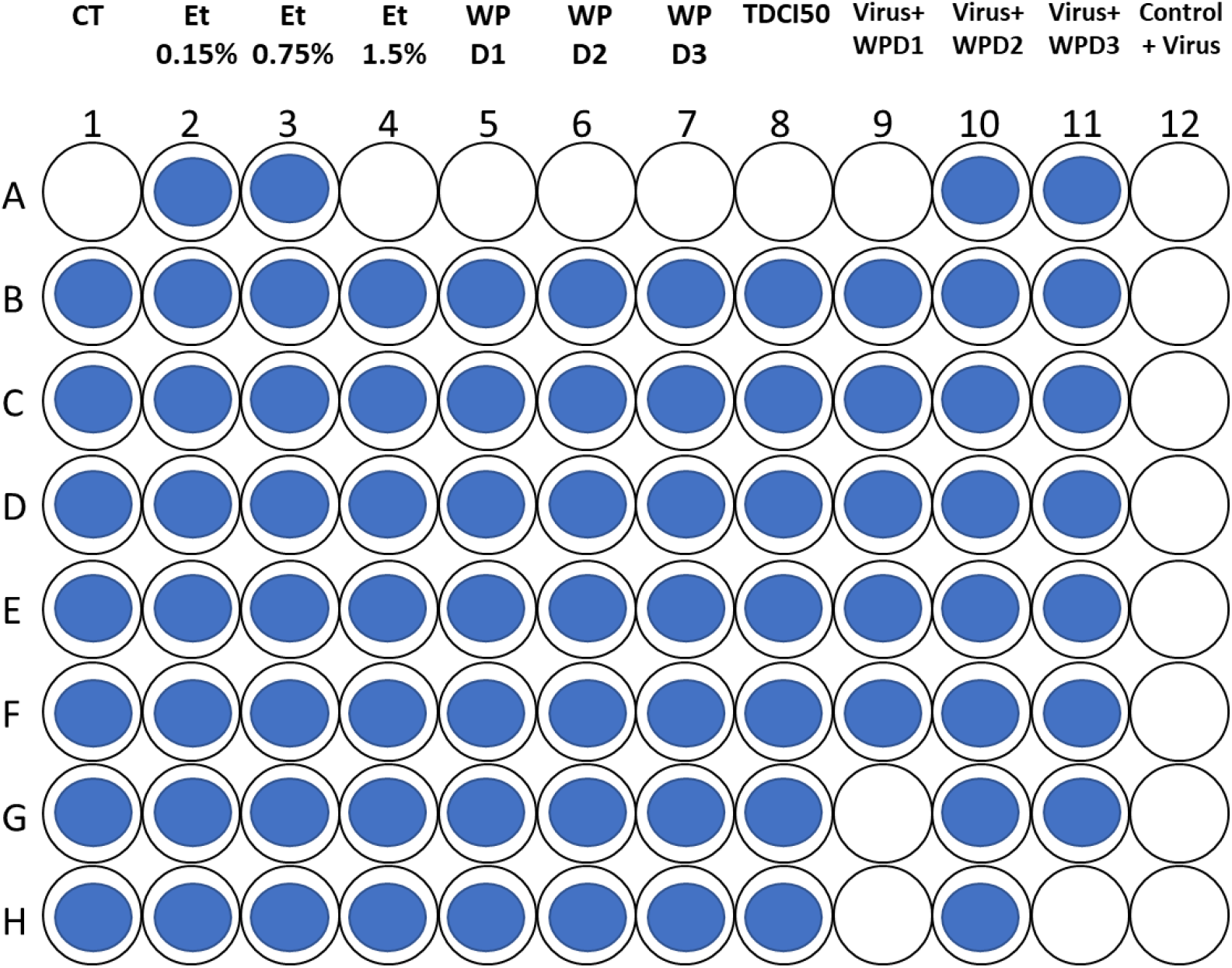
Cytotoxicity assay graphic representation. Shown data were collected at 27/07/2020 (first replicate). Columns are divided as indicated in experimental design. Blue-coloured dots represents viable cells. White-coloured dots represents unviable cells (dead). Note this replica had a TCID50 control weak (not kill 50% of Vero E6 cells). This example is shown to observe the marked cytoprotective effect in WPD2 and WPD3 columns comparing to 10-fold viral load positive control (100% of cellular death). No dose related WP D1, D2 and D3 cytotoxicity was observed on epithelial-derived Vero E6 cells (ICEP4 raw material, columns 5, 6 and 7).

Marked significant differences were found between groups for at least one group as stated by ANOVA of a factor (ICEP4 treatment) (Table 2). No differences were found by analysing the effect of solvents or raw material vs Vero culture in a two-sample t-test assuming equal variances (Table 3).

**Table 2.** Analysis of variance of a factor (ICEP4 treatment) of *in vitro* challenge test. Null hypothesis was considered as no observed differences between groups. *P < 0.05, **P < 0.01 and ***P < 0.001. P < 0.05 was considered as significant.

| *Origin of variations* | *F* | *Probability* | *Critical value for F* |
| --- | --- | --- | --- |
| Between groups | 29.03349282 | 0.000*** | 2.216308646 |

**Table 3.**
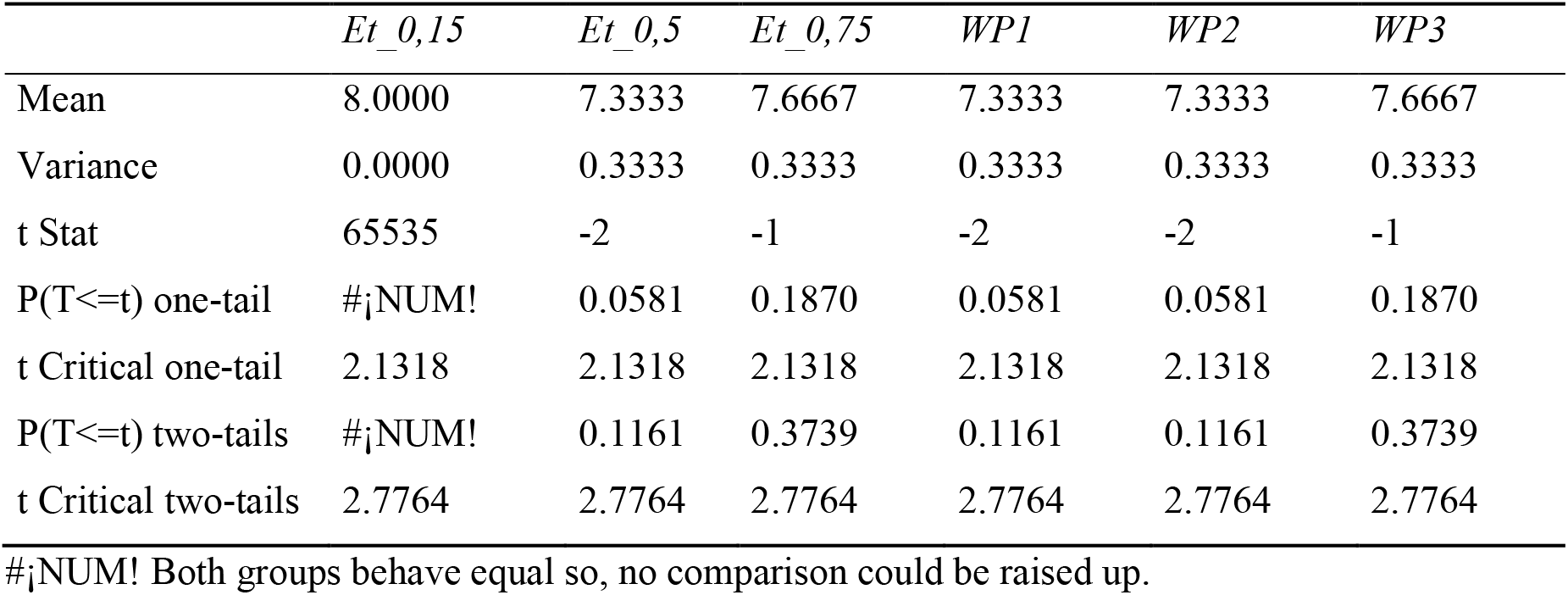
Two-sample t-test assuming equal variances results of *in vitro* challenge test for solvents and raw material. Non-treated Vero E6 2% medium culture (CT) group was considered treatment-negative control (maximum cell viability) for comparison (data no shown). Ethanol 0.15%, 0.75% and 1,5% respectively were considered as solvent control. ICEP4 doses were 6.9 μM (WP1), 34.5 μM (WP2) and 69 μM (WP3) and considered as ICEP4-treated non-SARS-CoV-2-challenged groups. *P < 0.05, **P < 0.01 and ***P < 0.001. P < 0.05 was considered as significant.

Marked increases in cell viability were observed when comparing to TCDI50 control were found in SARS-CoV-2-infected ICEP4-treated groups, corresponding to 34.5 μM (WP2) and 69 μM (WP3) doses (Table 4). Significant differences were also found between these groups when analysing two-sample t-test results, assuming equal variances, thus confirming preliminary descriptive results. Interpreting these results, ANOVA significance can be directly correlated to cytoprotective effects of ICEP4 treatment.

**Table 4.** Two-sample t-test assuming equal variances results of *in vitro* challenge test for SARS-CoV-2-infected ICEP4-treated groups. TCID50 SARS-CoV-2-infected cell death-positive group was considered for comparison (data no shown). *P < 0.05, **P < 0.01 and ***P < 0.001. P < 0.05 was considered as significant. ICEP4 doses were 6.9 μM (WP1), 34.5 μM (WP2) and 69 μM (WP3). Tissue Culture Infectious Dose 50% (TCID50) of SARS-CoV-2 (SARS) was determined in 1.47 × 10^6^/mL.

|  | SARS+WP1 | SARS+WP2 | SARS+WP3 |
| --- | --- | --- | --- |
| Mean | 7 | 7.667 | 7.667 |
| Variance | 1 | 0.333 | 0.333 |
| t Stat | 1.732 | 2.530 | 2.530 |
| P(T<=t) one-tail | 0.079 | 0.032* | 0.032* |
| t Critical one-tail | 2.132 | 2.132 | 2.132 |
| P(T<=t) two-tails | 0.158 | 0.065 | 0.065 |
| t Critical two-tails | 2.776 | 2.776 | 2.776 |

### 3.2. Cytotoxicity on B and T lymphocytes, NK and NKT cells and macrophages

No toxicity on CD4 and CD8 T cells, B cells, or NKT was seen at the tested does of ICEP4 (Tables 5 and 6; Figure 3). However, increased Annexin-V staining, indicating slight decreases in cellular viability, were observed in NK cells at the 100 μM dose. A similar effect was also found in macrophages, which showed a 30% reduction in cell viability at 100 μM of ICEP4 (Figure 3).

**Table 5.**
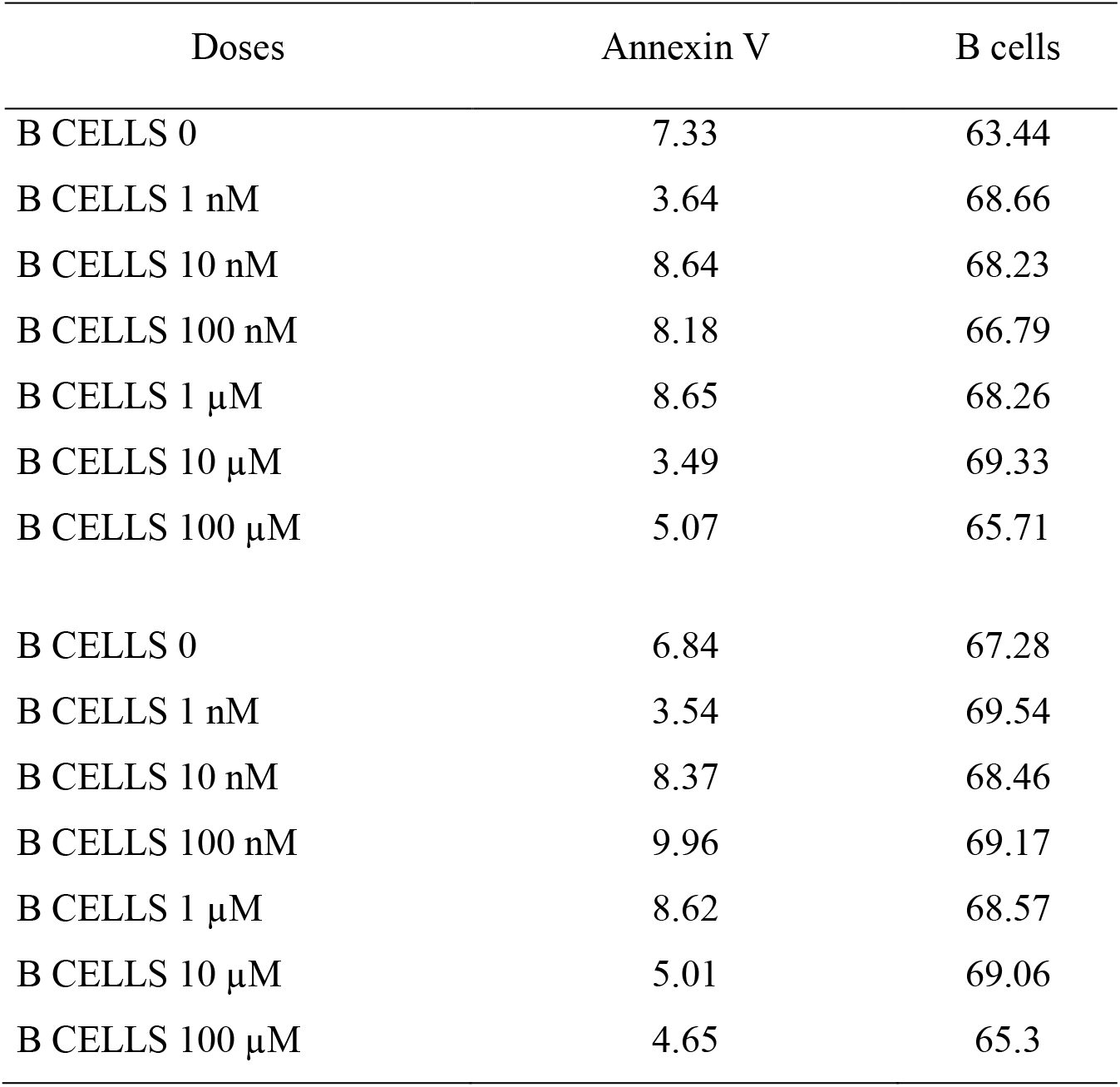
Annexin V staining (%) of B lymphocytes after ICEP4 challenge.

| Doses | Annexin V | B cells |
| --- | --- | --- |
| B CELLS 0 | 7.33 | 63.44 |
| B CELLS 1 nM | 3.64 | 68.66 |
| B CELLS 10 nM | 8.64 | 68.23 |
| B CELLS 100 nM | 8.18 | 66.79 |
| B CELLS 1 μM | 8.65 | 68.26 |
| B CELLS 10 μM | 3.49 | 69.33 |
| B CELLS 100 μM | 5.07 | 65.71 |
| B CELLS 0 | 6.84 | 67.28 |
| B CELLS 1 nM | 3.54 | 69.54 |
| B CELLS 10 nM | 8.37 | 68.46 |
| B CELLS 100 nM | 9.96 | 69.17 |
| B CELLS 1 μM | 8.62 | 68.57 |
| B CELLS 10 μM | 5.01 | 69.06 |
| B CELLS 100 μM | 4.65 | 65.3 |

**Table 6.** Annexin V staining (%) on immune cells (CD4+, CD8+, NK+ and NKT+ cells), and CD4+, CD8+, NK+ and NKT+ staining of lymphocytes T and NK after ICEP4 challenge.

| Data set | Annexin V |  |  |  | Lymphocyte marker |  |  |  |
| --- | --- | --- | --- | --- | --- | --- | --- | --- |
|  | CD4+ | CD8+ | NK Cells | NKT Cells | CD4+ | CD8+ | NK | NKT |
| T NK CELLS 000 00063130.715 | 17.66 | 7.25 | 5.6 | 10.12 | 4.09 | 9.09 | 7.28 | 18.37 |
| T NK CELLS 001 00063137.722 | 17.56 | 2.55 | 6.4 | 4.52 | 5.21 | 7.33 | 10.34 | 16.55 |
| T NK CELLS 010 00063136.721 | 14.02 | 2.66 | 4.22 | 4.13 | 4.4 | 8.83 | 8.72 | 17.77 |
| T NK CELLS 100 00063135.720 | 13.76 | 2.96 | 3.74 | 4.71 | 4.3 | 9.1 | 8.95 | 17.83 |
| T NK CELLS 001 00063134.719 | 14.21 | 2.22 | 3.86 | 4.41 | 4.44 | 8.87 | 8.62 | 17.72 |
| T NK CELLS 010 00063133.718 | 14.29 | 1.21 | 6.45 | 2.16 | 6.9 | 6.01 | 7.24 | 16.89 |
| T NK CELLS 100 00063132.717 | 20.47 | 0.76 | 11.42 | 0.63 | 4.41 | 6.25 | 8.33 | 18.68 |
| T NK CELLS 000 00063139.724 | 15.79 | 5.07 | 5.29 | 7.33 | 5.07 | 8.51 | 6.1 | 16.41 |
| T NK CELLS 001 00063146.731 | 15.72 | 0.26 | 5.52 | 1.77 | 6.68 | 6.11 | 8.63 | 16.46 |
| T NK CELLS 010 00063145.730 | 12.92 | 2.37 | 4.34 | 3.98 | 4.45 | 8.64 | 8.26 | 17 |
| T NK CELLS 100 00063144.729 | 12.47 | 2.46 | 3.79 | 5.24 | 3.87 | 8.69 | 7.35 | 17.54 |
| T NK CELLS 001 00063143.728 | 11.45 | 2.44 | 2.65 | 4.72 | 4.24 | 8.38 | 8.44 | 17.63 |
| T NK CELLS 010 00063142.727 | 15.04 | 0.67 | 5.82 | 1.85 | 6.31 | 6.27 | 6.23 | 16.65 |
| T NK CELLS 100 00063141.726 | 12.08 | 0.19 | 10.51 | 0.65 | 9.85 | 4.86 | 8.18 | 18.51 |

**Figure 3.**
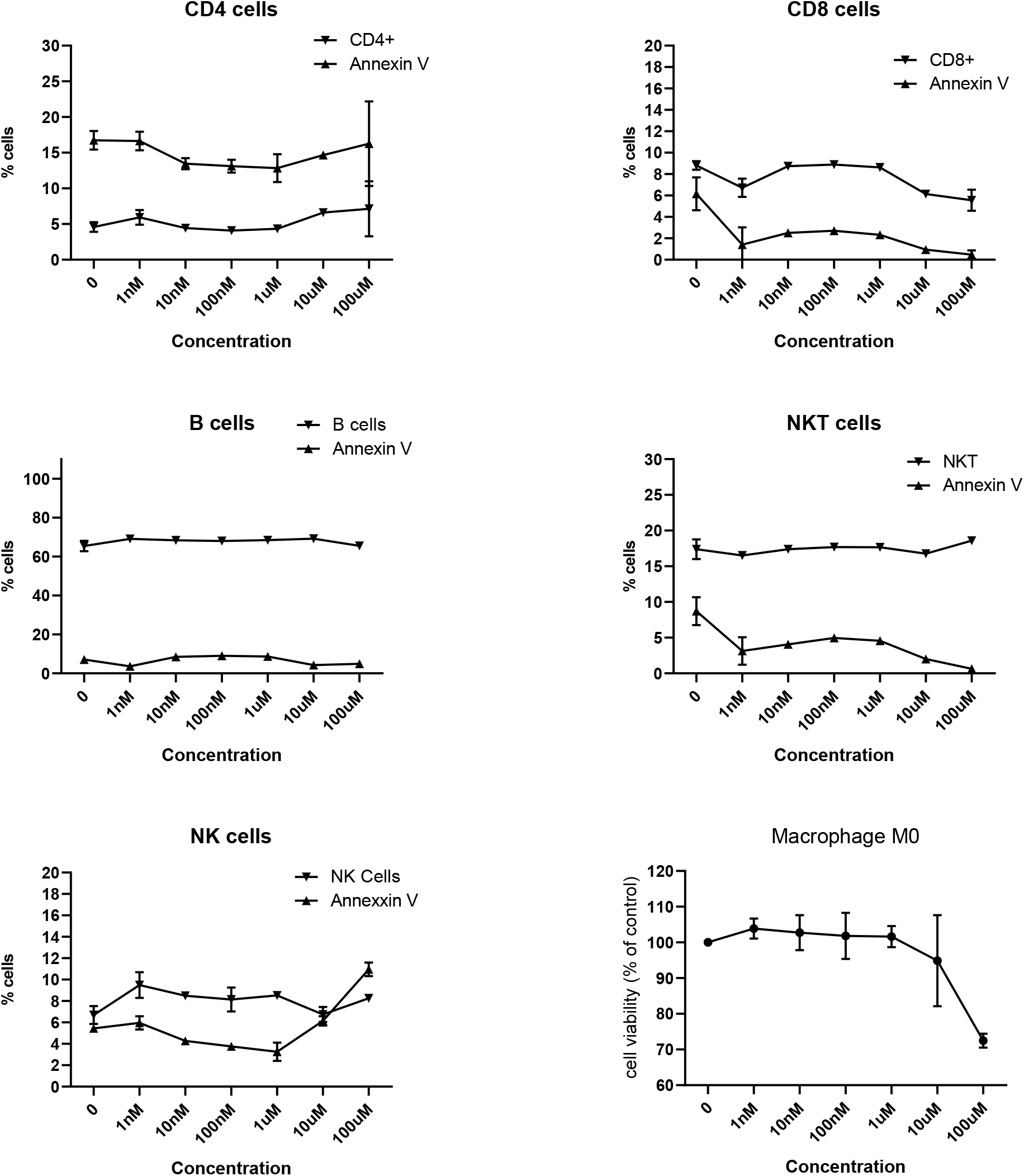
Effect of ICEP4 on cellular viability of M0 macrophages, B, CD4^+^ T and CD8^+^ T lymphocytes, Natural Killer (NK) and Natural Killer T (NKT) cells. Bone marrow derived macrophages (M0) or mouse splenocytes were incubated with different compound concentrations for 24h. Subsequently, the percentage of viable cells was determined using antibodies specific for T, B, NK and NKT cells and Annexin V staining as indicated in methods section. For M0 macrophages cell viability was measured by Presto Blue^TM^ assay. Individual points represent the mean value ±SD of 3 independent experiments.

Remarkably, ICEP4 plant-derived extract did not induce an inflammatory response in M0 macrophages at any of the tested doses, which we were able to verify via the release of IL6 and TNFα after challenge (Table 7). IL6 and TNFα values were very low and, in some samples, very close to or even below the limit of detection (detection limit IL6: 4 pg/mL, TNFα: 8 pg/mL). As expected, LPS induced a high inflammatory response thereby confirming macrophage functionality.

**Table 7.** IL6 and TNFα analyses from M0 macrophages supernatant.

| | | IL6 | | media | TNFα | | media |
| --- | --- | --- | --- | --- | --- | --- | --- |
| **M1** | control- | 4.8 | 5.52 | 5.16 pg/ml | 20.52 | 28.96 | 24.74 pg/ml |
| **M2** | 0 | 4.48 | 5.28 | 4.88 pg/ml | 17.08 | 28.32 | 22.7 pg/ml |
| **M3** | 1nM | 4.6 | 5.28 | 4.94 pg/ml | 19.16 | 25.84 | 22.5 pg/ml |
| **M4** | 10nM | 4.4 | 5.8 | 5.1 pg/ml | 17.48 | 32.52 | 25 pg/ml |
| **M5** | 100nM | 4.48 | 6.96 | 5.72 pg/ml | 21 | 38.2 | 29.6 pg/ml |
| **M6** | 1uM | 5.04 | 0.12 | 2.58 pg/ml | 24.68 | 0.04 | 12.36 pg/ml |
| **M7** | 10uM | 7.12 | 0.16 | 3.64 pg/ml | 29.64 | 0.76 | 15.2 pg/ml |
| **M8** | 100uM | 5.16 | 0.24 | 2.7 pg/ml | 26.44 | 0 | 13.22 pg/ml |
| **M9** | Diluyente | 5.52 | 0.12 | 2.82 pg/ml | 27.04 | 0.36 | 13.7 pg/ml |
| **Control +** | LPS | 605.44 pg/ml |  |  | 1236.12 pg/ml |  |  |

### 3.3. *In vivo* toxicity of ICEP4

Twenty-four hr acute and 21-day chronic *in vivo* toxicity studies were performed on ICEP4 in *C. elegans* (Figure 4). No toxicity was observed from doses of 1 nM to 100 μM in either assay. In both assays, only the highest dose showed toxicity (1 mM), which is presumably due to the higher ethanol concentration (20%) and not to the active ingredient, ICEP4, as confirmed by the SHAM control (Figure 4).

**Figure 4.**
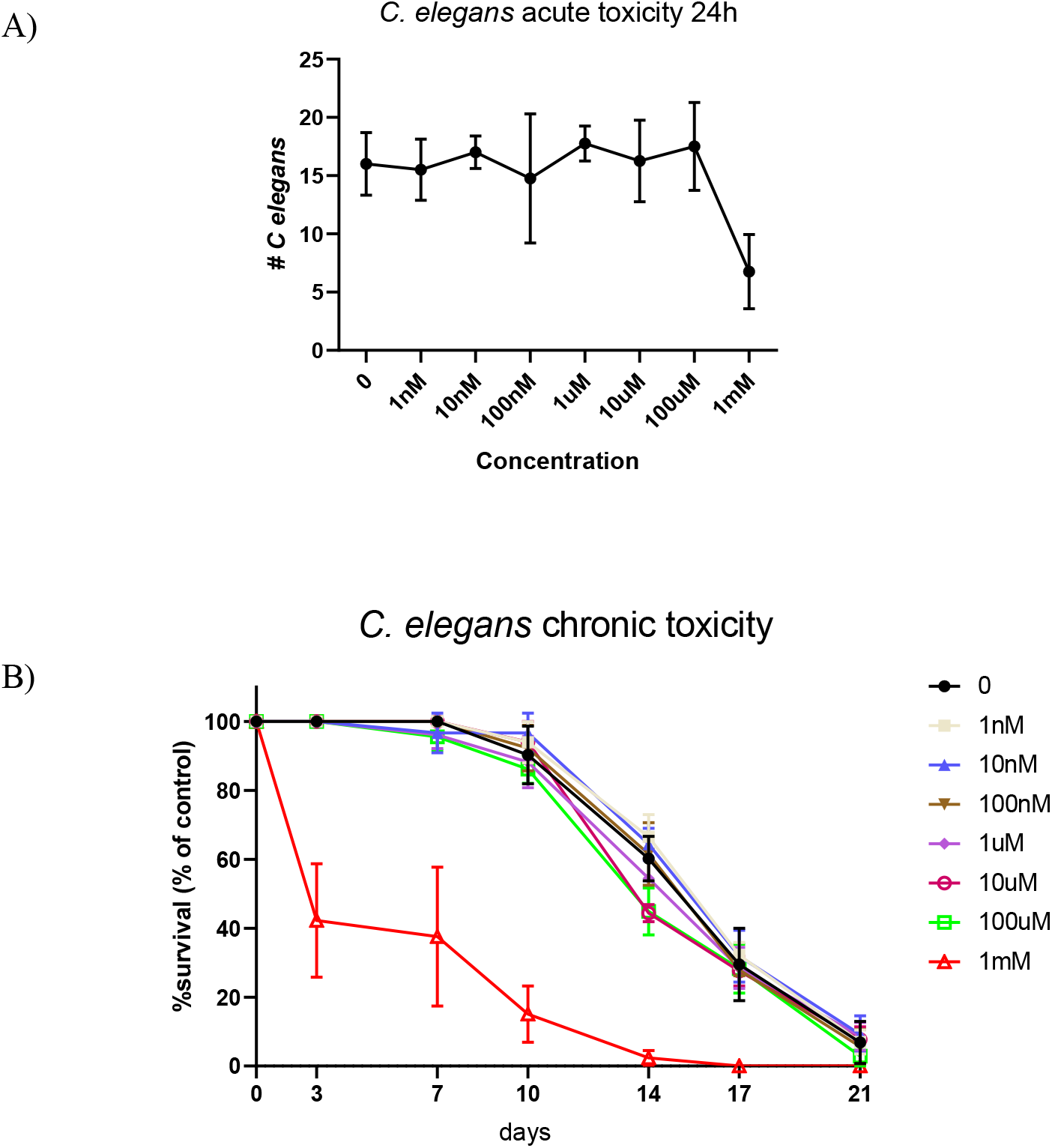
Acute (A) and chronic (B) toxicity of ICEP4 on *C. elegans* survival. Worms were incubated with different compound concentrations for 24h or 21 days and the percentage of viable worms was determined as indicated in methods. Individual points represent the mean value ±SD of 3 independent experiments.

## 4. Discussion

Antiviral herbal drugs have been widely used on the clinical frontline against respiratory diseases. Traditional Chinese medicines (TCM), Ayurveda medicine (AM), and European herbal drugs (EHD) are highly encouraged as adjuvant therapies in COVID19, supported by historically well documented efficacy studies against many viral infections, including influenza, SARS, and MERS (16, 26, 27). Vegetal drugs are frequently used in herbal decoctions for lung clearing and detoxification in the clinical mitigant treatment of respiratory diseases and, recently, also COVID19, with the most common being *Astragalus membranaceus*, *Glycyrrhizae uralensis*, *Saposhnikoviae divaricata*, *Rhizoma Atractylodis*, *Macrocephalae*, *Lonicerae Japonicae Flos*, *Fructus forsythia*, *Atractylodis Rhizoma*, *Radix platycodonis*, *Agastache rugosa*, *Cyrtomium fortune* J. Sm., *Withania somnifera* (Ashwagandha), *Tinospora cordifolia* (Guduchi), *Asparagus racemosus* (Shatavari), *Phylanthus embelica* (Amalaki), and *Glyceriza glabra* (Yashtimadhu) (28, 29). As reviewed by Sarker and Nahal (2004), many species of the *Angelica* genus, e.g., *A. acutiloba*, *A. archangelica*, *A. atropupurea*, *A. dahurica*, *A. japonica*, *A. glauca*, *A. gigas*, *A. koreana*, *A. sinensis*, *A. sylvestris*, etc., have been used for centuries as anti-inflammatory, expectorant, and diaphoretic substances, as well as remedies for colds, flu, influenza, coughs, chronic bronchitis, pleurisy, headaches, fever, and diverse bacterial and fungal infections, among others (16). Active principles isolated from these plants mainly include various types of coumarins, acetylenic compounds, chalcones, sesquiterpenes, and polysaccharides (9, 10, 16). Frequently, most of the existing conventional antiviral treatments lead to the development of viral resistance in addition to the problems of side effects, viral re-emergence, and viral dormancy (2). Therefore, the WHO also supports and welcomes innovations around the world regarding scientifically proven traditional medicine, in order to increase the clinical alternatives of safe antiviral therapies (30).

*Angelica archangelica*-related ICEP4 has shown marked, significant *in vitro* cytoprotective effects in SARS-CoV-2-challenged Vero E6 cellular cultures at doses of 34.5 and 69 μM (0.75 and 1.5 μg/dose, respectively). Successful ICEP4 doses against SARS-CoV-2 are included within the maximum TTC of furocoumarin as a herbal preparation or remedy (24). Total daily human exposure to coumarins from dietary or cosmetic sources is 0.06 mg/Kg/day, with a total daily dose of 0.2% furocoumarins (1.2 μg/Kg/day) (31). No adverse effects of coumarin have been reported in susceptible species in response to doses which are more than 100-fold the maximum human daily intake. Successful ICEP4 doses used in these experiments are within the range of non-toxic use of furocoumarins (1.5 μg in 100 μL per well) (13, 18, 31). It is worth mentioning that the non-cytotoxic proof-of-concept ICEP4 dose is 4-fold lower than the antiproliferative cytotoxicity threshold observed in previous *Angelica*-related studies (32–34). These promising results might open the possibility to further studies for druggability of ICEP4. Remarkably, a highly-virulent strain of SARS-CoV-2 was used during these experiments, therefore, we can postulate that the ICEP4-related herbal drug is a promising potential treatment for COVID19.

EHD, AM and TCM, either alone or in combination, have been used for centuries in clinical and prophylactic antiviral treatments, and have proven their efficacy when subjected to rigorous scientific investigation (35). Additionally, these therapies have played a major role in the discovery and development of many antiviral drugs based on their structural moieties, with the classical examples being emetine and quinine (36). Naturally occurring scaffolds, such as coumarins, display a wide spectrum of pharmacological activities, including anticancer, antibiotic, and antidiabetic activities, among others, via action on multiple targets. In this view, various coumarin-based hybrids possessing diverse medicinal attributes have been synthesised in the past five years by conjugating a coumarin moiety with other therapeutic pharmacophores (37). Antiviral mechanistic studies of TCM, AM, and/or EHD have revealed how they interfere with the viral life cycle, as well as virus-specific host targets (4). Herbal drugs have demonstrated a wide range of antiviral mechanisms that include inhibition of 3CLpro protein (Chinese *Rhubarb* extracts, *Houttuynia chordata* extract, hesperetin, etc.), blocking or inhibition of the viral RNA-dependent RNA polymerase activity, and inhibition of inflammatory cytokines, such as TNFα, IL1β, and IL6 (*Fructus forsythiae*) (38). A psychoneuroimmune mechanism has been highlighted as a possible immune-mediated pathway in COVID19 treatment support (39, 40).

Prophylactic therapeutics with potential immunomodulatory activity have been postulated as add-on treatments for COVID19 (29). Many medicinal compounds and natural products have exhibited several antiviral mechanisms that prevent early stages of infection, including viral attachment and penetration (36, 41). Psoralens, the main moiety of furocoumarins, may react directly with pyrimidine nucleotides to form mono- and di-adducts in DNA of even interstrand cross links (6). Another route of psoralen toxicity derives from the ability of UV-A photoactivated furanocoumarins to react with grand state oxygen, generating toxic oxyradicals capable of inactivating proteins within cells (6, 7). Due to this reactivity, a broad range of therapeutic applications requiring inhibitors of cell division (main drug targets are the cytochrome P450 superfamily) have been suggested, such as vitiligo, psoriasis, and several type of cancers, including T cell lymphoma (7, 32–34). Taking together the data from the cytotoxicity and *in vivo* assays, we postulate that furocoumarin-derived ICEP4 has shown very little, or negligible evidence of toxicity towards immune and epithelial-derived cells. NK and macrophages showed slight decreases in viability when exposed to the highest dose, probably due to the same effect of solvent stated for the *in vivo* assays. It is worth mentioning that *Angelica*-based furocoumarin, the gold standard, is phototoxic and affects cellular viability within the studied doses (32, 34). Several structural changes, well-established compound concentration and composition differences are expected by meaning of geographic, stational or plant-related issues, many of them can be used as drug template design (27). Therefore, it is postulated that ICEP4, despite being extracted and purified in the same manner as other *Angelica*-related furocoumarins, possesses differences in chemical or racemic compounding composition relating to the extraction method, which leads to marked differences in terms of toxicity and antiviral efficacy (SARS-CoV-2 infection model). These facts may let us assume that an unknown different furocoumarin-related compound has been discovered or, at least, first observed. The extraction method of patent-pending ICEP4 has showed previously strong manufacturing robustness, long-lasting stability, and repeated chemical consistency. Nonetheless, additional chemical-related data regarding ICEP4 are urgently needed, in order to highlight several concerns regarding its use as a herbal drug.

TCM, AM, and EHD-prescribed herbal drugs decrease the severity and mortality rate of COVID19 (28, 35, 38, 41). Several drugs, such as *Nigella sativa*, natural honey, artemisinin, curcumin, *Boswellia*, and vitamin C, are in a clinical trials for COVID19 treatment (35). Although promising clinical evidence in the treatment of a diverse range of respiratory infections is available for *Angelica archangelica* (16, 31), randomised human clinical trials are required to evaluate the efficacy and safety of ICEP4 in COVID19 patients. Historical clinical evidence of *Angelica*-related remedies in the treatment of respiratory syndromes, its safe toxicity profile, and the appropriate range of therapeutic doses may lead to the use of ICEP4 as a prophylactic mucosal-related non-systemic anti-SARS-CoV-2 therapy. In this view, and encouraged by WHO demands, ICEP4 herbal drug may be suitable for complete drug development under the COVID19 regulatory pathway.

## 5. Acknowledgment

Authors wish to thank Eduardo Pérez and Dr. Javier Agulló for their kind support in reviewing this manuscript for patentability issues. Authors wish to thank Eugenia Puentes (Biofabri, Porriño, Spain) for providing Vero E6 cells. Proof-Reading-Services.com Ltd was contracted to perform an idiomatic review (British English, Vancouver referencing).

## Author contributions

Original ICEP4 batch and stability data (Ezio Panzeri). Data collection, samples management and analytic ICEP4 control by HPLC, flow cytometry and staining techniques (Adriana Toledo Núñez, María Celaya Fernández, Iratxe Uranga-Murillo, Maykel Arias Cabrero and Ariel Ramírez Labrada). Study design and data interpretation (Iván José Galindo-Cardiel, Ariel Ramirez Labrada). Writing and editing tables and figures (Iván José Galindo-Cardiel, Ariel Ramírez Labrada, Ezio Panzeri). Quality assurance (Iván José Galindo-Cardiel, Julián Pardo).

## Disclosure

Dr. Iván José Galindo-Cardiel and Ezio Panzeri are co-authors of ICEP4-related patent application (GB2017123.7).

**Funding**

## Notes

i Iratxe Uranga-Murillo is supported by a predoctoral contract from Aragón Government.

ii Dr. Maykel Arias is supported by a Juan de la Cierva postdoctoral contract.

iii Dr. Julian Pardo’s laboratory is funded by FEDER (Fondo Europeo de Desarrollo Regional, Gobierno de Aragón -Group B29_17R-, Ministerio de Ciencia, Innovación e Universidades - MCNU-, Agencia Estatal de Investigación -SAF2017‐83120‐C2‐1‐R-), Instituto de Salud Carlos III, Fundación Inocente Inocente, ASPANOA and Carrera de la Mujer de Monzón.

### Competing Interest Statement

Dr. Iván José Galindo-Cardiel and Ezio Panzeri are co-authors of ICEP4-related patent application (GB2017123.7).
Iratxe Uranga-Murillo is supported by a predoctoral contract from Aragón Government.
Dr. Maykel Arias is supported by a Juan de la Cierva postdoctoral contract.
Dr. Julian Pardo’s laboratory is funded by FEDER (Fondo Europeo de Desarrollo Regional, Gobierno de Aragón -Group B29_17R-, Ministerio de Ciencia, Innovación e Universidades -MCNU-, Agencia Estatal de Investigación -SAF2017‐83120‐C2‐1‐R-), Instituto de Salud Carlos III, Fundacion Inocente Inocente, ASPANOA and Carrera de la Mujer de Monzón.

### Summary of Updates

British English - Idiomatic revision

## References

1. Gorbalenya AE, Baker SC, Baric RS, de Groot RJ, Drosten C, Gulyaeva AA, et al. The species Severe acute respiratory syndrome-related coronavirus: classifying 2019-nCoV and naming it SARS-CoV-2. Nature Microbiology. 2020;5(4):536–44.

2. Kumari P, Rawat K, Saha L. Pipeline Pharmacological Therapies in Clinical Trial for COVID-19 Pandemic: a Recent Update. Current pharmacology reports. 2020:1–13.

3. Alvarez A, Cabia L, Trigo C, Bandres AC, Bestue M. Prescription profile in patients with SARS-CoV-2 infection hospitalised in Aragon, Spain. European journal of hospital pharmacy: science and practice. 2020.

4. Boukhatem MN, Setzer WN. Aromatic Herbs, Medicinal Plant-Derived Essential Oils, and Phytochemical Extracts as Potential Therapies for Coronaviruses: Future Perspectives. Plants. 2020;9(6).

5. Egan DA, O'Kennedy R. Rapid and sensitive determination of coumarin and 7-hydroxycoumarin and its glucuronide conjugate in urine and plasma by high-performance liquid chromatography. Journal of chromatography. 1992;582(1-2):137–43.

6. Finn GJ, Kenealy E, Creaven BS, Egan DA. *In vitro* cytotoxic potential and mechanism of action of selected coumarins, using human renal cell lines. Cancer letters. 2002;183(1):61–8.

7. Egan D, O'Kennedy R, Moran E, Cox D, Prosser E, Thornes RD. The pharmacology, metabolism, analysis, and applications of coumarin and coumarin-related compounds. Drug metabolism reviews. 1990;22(5):503–29.

8. Lake BG. Coumarin metabolism, toxicity and carcinogenicity: relevance for human risk assessment. Food and chemical toxicology: an international journal published for the British Industrial Biological Research Association. 1999;37(4):423–53.

9. Keating G, O’Kennedy R. The Chemistry and Occurrence of Coumarins. Coumarins: Biology, Applications and Mode of Action. 1997.

10. Pereira TM, Franco DP, Vitorio F, Kummerle AE. Coumarin Compounds in Medicinal Chemistry: Some Important Examples from the Last Years. Current topics in medicinal chemistry. 2018;18(2):124–48.

11. Feng T, Downie SR, Yu Y, Zhang X, Chen W, He X, et al. Molecular systematics of *Angelica* and allied genera (*Apiaceae*) from the Hengduan Mountains of China based on nrDNA ITS sequences: phylogenetic affinities and biogeographic implications. Journal of plant research. 2009;122(4):403–14.

12. Karnick CR. Pharmacopoeial standards of herbal plants Delhi, India: Sri Satguru Publications; 1994. Available from: http://books.google.com/books?id=lwhtAAAAMAAJ.

13. (BAnz) B. Monographien der Kommission E. In: (Zulassungs-und Aufbereitungskommission am BGA für den humanmed. Bereich pTuS, editor. 1998.

14. Braun R, Norden‐Ehlert E, Surmann P, Wendt R, Wichtl M. Standardzulassungen für Fertigarzneimittel—Text and Kommentar. Stuttgart: Deutscher Apotheker Verlag. 1997.

15. Taufel A. A. Y. Leung and S. Foster: Encyclopedia of Common Natural Ingreadients Used in Food, Drugs and Cosmetics. Second edition, 649 pages. John Wiley & Sons, Inc., New York, Chichester, Brisbane, Toronto, Singapore 1996. Price: 115.00 £. Food / Nahrung. 1996;40(5):289-.

16. Sarker SD, Nahar L. Natural medicine: the genus *Angelica*. Current medicinal chemistry. 2004;11(11):1479–500.

17. Kim SH, Lee SW, Park HJ, Lee SH, Im WK, Kim YD, et al. Anti-cancer activity of *Angelica gigas* by increasing immune response and stimulating natural killer and natural killer T cells. BMC complementary and alternative medicine. 2018;18(1):218.

18. Thanh PN, Jin W, Song G, Bae K, Kang SS. Cytotoxic coumarins from the root of *Angelica dahurica*. Archives of pharmacal research. 2004;27(12):1211–5.

19. Shin S, Joo SS, Park D, Jeon JH, Kim TK, Kim JS, et al. Ethanol extract of *Angelica gigas* inhibits croton oil-induced inflammation by suppressing the cyclooxygenase - prostaglandin pathway. Journal of veterinary science. 2010;11(1):43–50.

20. Widelski J, Popova M, Graikou K, Glowniak K, Chinou I. Coumarins from *Angelica lucida* L.--antibacterial activities. Molecules. 2009;14(8):2729–34.

21. Wedge DE, Klun JA, Tabanca N, Demirci B, Ozek T, Baser KH, et al. Bioactivity-guided fractionation and GC/MS fingerprinting of *Angelica sinensis* and *Angelica archangelica* root components for antifungal and mosquito deterrent activity. Journal of agricultural and food chemistry. 2009;57(2):464–70.

22. Kang SY, Kim YC. Neuroprotective coumarins from the root of *Angelica gigas*: structure-activity relationships. Archives of pharmacal research. 2007;30(11):1368–73.

23. Deng S, Chen SN, Yao P, Nikolic D, van Breemen RB, Bolton JL, et al. Serotonergic activity-guided phytochemical investigation of the roots of *Angelica sinensis*. Journal of natural products. 2006;69(4):536–41.

24. (HMPC) COHMP. Risks associated with furocoumarins contained in preparations of *Angelica archangelica* L. In: (EMA) EMA, editor. Post-authorisation Evaluation of Medicines for Human Use. London 2007. p. 1–22.

25. Porta-de-la-Riva M, Fontrodona L, Villanueva A, Cerón J. Basic *Caenorhabditis elegans* methods: synchronization and observation. Journal of visualized experiments: JoVE. 2012(64):e4019.

26. Leung PC. The efficacy of Chinese medicine for SARS: a review of Chinese publications after the crisis. The American journal of Chinese medicine. 2007;35(4):575–81.

27. Islam MT, Sarkar C, El-Kersh DM, Jamaddar S, Uddin SJ, Shilpi JA, et al. Natural products and their derivatives against coronavirus: A review of the non-clinical and pre-clinical data. Phytotherapy research : PTR. 2020;34(10):2471–92.

28. Huang F, Li Y, Leung EL, Liu X, Liu K, Wang Q, et al. A review of therapeutic agents and Chinese herbal medicines against SARS-COV-2 (COVID-19). Pharmacological research. 2020;158:104929.

29. Balasubramani SP, Venkatasubramanian P, Kukkupuni SK, Patwardhan B. Plant-based Rasayana drugs from Ayurveda. Chinese journal of integrative medicine. 2011;17(2):88–94.

30. World Health O. WHO traditional medicine strategy: 2014-2023. Geneva: World Health Organization; 2013 2013.

31. Council AB. Angelica root: Integrative Medicine Communications 2000. Available from: http://cms.herbalgram.org/expandedE/Angelicaroot.html?ts=1606421458&signature=48b9cd2ad7fcc5d691784102f498e149&ts=1606558508&signature=2a594d9c3c4a6b38629367878c8d059a.

32. Sigurdsson S, Ogmundsdottir HM, Gudbjarnason S. The cytotoxic effect of two chemotypes of essential oils from the fruits of *Angelica archangelica* L. Anticancer research. 2005;25(3B):1877–80.

33. Sigurdsson S, Ogmundsdottir HM, Hallgrimsson J, Gudbjarnason S. Antitumour activity of *Angelica archangelica* leaf extract. In vivo. 2005;19(1):191–4.

34. Sigurdsson S, Ogmundsdottir HM, Gudbjarnason S. Antiproliferative effect of *Angelica archangelica* fruits. Zeitschrift fur Naturforschung C, Journal of biosciences. 2004;59(7-8):523–7.

35. Interventional studies for COVID-19 [Internet]. 2020 [cited 28 Nov 2020]. Available from: https://clinicaltrials.gov/ct2/results?cond=COVID-19.

36. Mani JS, Johnson JB, Steel JC, Broszczak DA, Neilsen PM, Walsh KB, et al. Natural product-derived phytochemicals as potential agents against coronaviruses: A review. Virus research. 2020;284:197989.

37. Singh H, Singh JV, Bhagat K, Gulati HK, Sanduja M, Kumar N, et al. Rational approaches, design strategies, structure activity relationship and mechanistic insights for therapeutic coumarin hybrids. Bioorganic & medicinal chemistry. 2019;27(16):3477–510.

38. Yang Y, Islam MS, Wang J, Li Y, Chen X. Traditional Chinese Medicine in the Treatment of Patients Infected with 2019-New Coronavirus (SARS-CoV-2): A Review and Perspective. International journal of biological sciences. 2020;16(10):1708–17.

39. Rajkumar RP. Ayurveda and COVID-19: Where psychoneuroimmunology and the meaning response meet. Brain, behavior, and immunity. 2020;87:8–9.

40. Sigurdsson S, Gudbjarnason S. Inhibition of acetylcholinesterase by extracts and constituents from *Angelica archangelica* and Geranium sylvaticum. Zeitschrift fur Naturforschung C, Journal of biosciences. 2007;62(9-10):689–93.

41. Gyawali R, Paudel PN, Basyal D, Stezer WN, Lamichhane S, Kumar M. A Review on Ayurvedic Medicinal Herbs as Remedial Perspective for COVID-19. Journal of Karnali Academy of Health Sciences. 2020;3.

